# Abundant Aβ fibrils in ultracentrifugal supernatants of aqueous extracts from Alzheimer’s disease brains

**DOI:** 10.1101/2022.10.18.512754

**Authors:** Andrew M. Stern, Yang Yang, Angela L. Meunier, Wen Liu, Yuqi Cai, Maria Ericsson, Lei Liu, Michel Goedert, Sjors H. W. Scheres, Dennis J. Selkoe

**Affiliations:** Ann Romney Center for Neurologic Diseases, Department of Neurology, Harvard Medical School and Brigham and Women’s Hospital, Boston, MA 02115; Medical Research Council Laboratory of Molecular Biology, Cambridge UK; Harvard Medical School Electron Microscopy Facility, Boston, USA

**Keywords:** Aβ, Alzheimer’s disease, fibrils, oligomers, therapeutic antibodies, amyloid

## Abstract

Soluble aggregates of amyloid-β (Aβ), often called oligomers, are believed to be principal drivers of neurotoxicity, spreading of pathology, and symptoms in Alzheimer’s disease (AD), but little is known about their structures in human brain. Aβ oligomers have been defined as aggregates found in supernatants following ultracentrifugation of aqueous extracts. We now report the unexpected presence of abundant Aβ fibrils in high-speed supernatants from AD brains that were extracted by soaking in aqueous buffer. The fibrils did not appear to form during extract preparation, and their numbers by EM correlated with ELISA quantification of aggregated Aβ_42_. Cryo-EM structures of Aβ fibrils from aqueous extracts were identical to those from sarkosyl-insoluble AD brain homogenates. The fibrils in aqueous extracts were immunolabeled by lecanemab, an Aβ aggregate-directed antibody reported to improve cognitive outcomes in AD. We conclude that Aβ fibrils are abundant in aqueous extracts from AD brains and have the same structures as those from amyloid plaques. These findings have implications for understanding the nature of Aβ oligomers and for designing oligomer-preferring therapeutic antibodies.

## Introduction

Amyloid β (Aβ) and tau proteins convert from soluble monomers into insoluble fibrils when they aggregate. Interest in fibril formation derives from their presence in amyloid plaques and tau tangles in Alzheimer’s disease (AD) brains, and from the causation of neurodegenerative disease by mutations that accelerate aggregation of Aβ and tau.^1,2^ Fibrils can be extracted from *postmortem* brains and pelleted by differential ultracentrifugation in the presence of detergents such as N-lauryl sarcosine (sarkosyl). Detergent-insoluble Aβ has been reported to be less toxic than Aβ from aqueous ultracentrifugal supernatants prepared without detergent.^3^ Aqueous supernatants contain putatively soluble Aβ assemblies that disrupt synaptic plasticity^3–5^ and neurite morphology,^6,7^ promote tau phosphorylation,^6,8,9^ decrease memory in rodent models^3^ and accelerate Aβ seeding.^10–13^ Because such assemblies are not pelleted by ultracentrifugation and are not monomeric, they have been termed “soluble” oligomers, and their neurotoxicity makes them attractive therapeutic targets.

The most promising antibodies in advanced trials for AD target aggregated Aβ.^14^ They include lecanemab (BAN2401), a humanized mouse monoclonal antibody reported to improve cognitive outcomes in a phase 3 trial^15,16^ and to bind preferentially to soluble Aβ aggregates over insoluble fibrils.^15^ Here, we use the term “assembly” or “aggregate” instead of “oligomer,” because *oligo* implies a *few* monomers, whereas assemblies in aqueous extracts may contain thousands of non-covalently bound monomers. We use the term “protofilament” to mean a single stack of monomers in a β-pleated sheet; in the case of Aβ fibrils, two protofilaments form one fibril.^16^

Synthetic Aβ can aggregate *in vitro* into assemblies of various sizes. However, their relevance for AD is uncertain because of the difficulty in extracting assemblies of known structure and solubility from human brain.^17^ The term “soluble” is operational because Aβ aggregates are found in aqueous supernatants after ultracentrifugation, but ultracentrifugation protocols vary and are often poorly described. Most aqueously extracted aggregates of Aβ from AD brains have been reported to be high molecular weight (HMW): they elute in or near the void volume of size exclusion chromatography (SEC) columns (≥500 kDa).^18,19^ Low MW aggregates isolated from AD brain, including covalent dimers,^20^ are particularly toxic,^3,19^ but disassembly of HMW aggregates *ex vivo* is often necessary to observe LMW aggregates and demonstrate their toxicity.^3,6,19^ The structure of any of these aggregates is unknown.

Dynamic non-covalent interactions render extraction of Aβ assemblies under non-denaturing conditions challenging. A recent method uses soaking of minced bits of AD cerebral cortex in Tris-buffered saline (TBS) rather than tissue homogenization, to avoid shearing plaques and other large Aβ aggregates; these soaking extracts exhibit similar neurotoxicity to whole brain homogenates.^21,22^ We previously reported a non-denaturing method to further enrich for Aβ aggregates from soaking extracts of AD cortex by immunoprecipitation with the calcium-sensitive Aβ antibody B24.^23^ We were surprised to observe Aβ fibrils in the B24 eluates of putatively soluble aqueous extracts.

We now report that both Aβ fibrils and tau paired helical filaments (PHFs) are present in aqueous soaking extracts, even in the absence of immunoprecipitation. With sufficient centrifugal force, virtually all Aβ aggregates could be pelleted. By electron cryo-microscopy (cryo-EM), these fibrils had the same structures as those from sarkosyl-insoluble homogenates of AD brains.^16^ Our results suggest that many HMW Aβ aggregates from aqueous extracts of AD brains are amyloid fibrils. These fibrils can be immunolabeled by lecanemab, a putatively oligomer-preferring Aβ antibody that has been reported to improve cognitive function in early AD.

## Results

### Aβ fibrils can be pelleted from aqueous extracts of Alzheimer’s disease brain

For immunoprecipitation using the calcium-requiring Aβ antibody B24,^23^ we started with aqueous extracts prepared by mincing and soaking multiple regions of AD cortex in TBS without homogenization (Table 1).^21^ We previously immunoprecipitated Aβ with B24 in the presence of calcium and eluted Aβ using only the chelator EGTA to avoid conditions which might denature Aβ aggregates.^23^ The resulting HMW Aβ aggregates from AD brains could then be re-pelleted in a benchtop centrifuge at 20,000 *g* in a 1.5-ml tube for 1 h (pelleting distance ~1 cm), with no detectable Aβ remaining in the supernatants by ELISA. Examining these pellets by EM revealed the presence short Aβ fibrils.^23^

**Table 1.** The soaking extract method and its modifications for control experiments in this report

| Step # | Steps in standard protocol | Modifications of standard protocol for the specified control experiments, and relevant figure number |
| --- | --- | --- |
| 1 | Frozen tissue thawed, grey matter dissected | **Fig S3I-L:** Fresh tissue |
| 2 | Tissue chopped using McIlwain Tissue Chopper set to 0.5 mm width |  |
| 3 | Tissue bits soaked in extraction buffer 30 min 4°C with nutation in 50-ml tube | **Fig S2B:** Tissue bits first rinsed with ~15 volumes extraction buffer before soaking |
| 4 | 2,000 g fixed angle (pelleting distance ~4 cm) 10 min 4°C, top 90% supernatant collected |  |
| 5 | 200,000 g in SW41Ti (pelleting distance ~9 cm) 110 min 4°C, top 90% of supernatant collected | **Fig S2A:** Top 90% of supernatant divided into top 2 ml and bottom ~7 ml<br>**Fig 1C-G:** 20,000 – 475,000 g in TLA100.3 (pelleting distance ~2 cm) for 110 min 4°C; top 90% of supernatant collected |
| 6 | Aliquot 1 ml into 1.5-ml Eppendorf LoBind tubes; store -80°C | **Fig S2E:** Aliquot into siliconized or BSA-coated tubes<br>**Fig S3C, H:** Store at 4°C<br>**Fig S3D:** Dilute serially before freezing<br>**Fig S3E-F:** Add various Aβ aggregation inhibitors before freezing<br>**Fig S3G:** Filter through aluminum oxide filters before freezing |
| 7 (re-centrifugation) | Thaw aliquot(s), 20,000 g fixed angle (pelleting distance ~1 cm) 60 min 4°C. Retain input, supernatant and pellet for ELISA and/or EM |  |

We now asked if Aβ fibrils could be pelleted and observed without immunoprecipitation. We spun 1 ml TBS soaking extracts (Table 1, Step #7), choosing arbitrarily the same 20,000 *g* protocol for 1 h. Soluble Aβ aggregates would not be expected to pellet. However, a substantial proportion of aggregated Aβ (*i*.*e*., that which was detectable by a monomer-preferring ELISA only after denaturation in 5 M GuHCl) was depleted from the supernatants and recovered in the pellets (Fig 1A, center). The natively monomeric Aβ in the original supernatants (*i*.*e*., not requiring GuHCl denaturation for detection by the monomer-preferring ELISA) was not depleted by the re-centrifugation, as expected (Fig 1A, right). Examining the re-centrifugation pellets by EM revealed amyloid fibrils that reacted with N-terminal anti-Aβ antibody D54D2 (Fig 1B). In aqueous extracts from AD brains, we also noticed the presence of fibrils resembling PHFs that stained with antibodies directed against the tau C-terminus and against phosphotau (Fig S1). In this report, we focus our analysis on Aβ fibrils. No fibrils were observed in the pellets from aqueous extracts of two control brains (C1 and C2), and only a single fibril was seen on an entire EM grid of a third control brain (C3).

**Fig 1.**
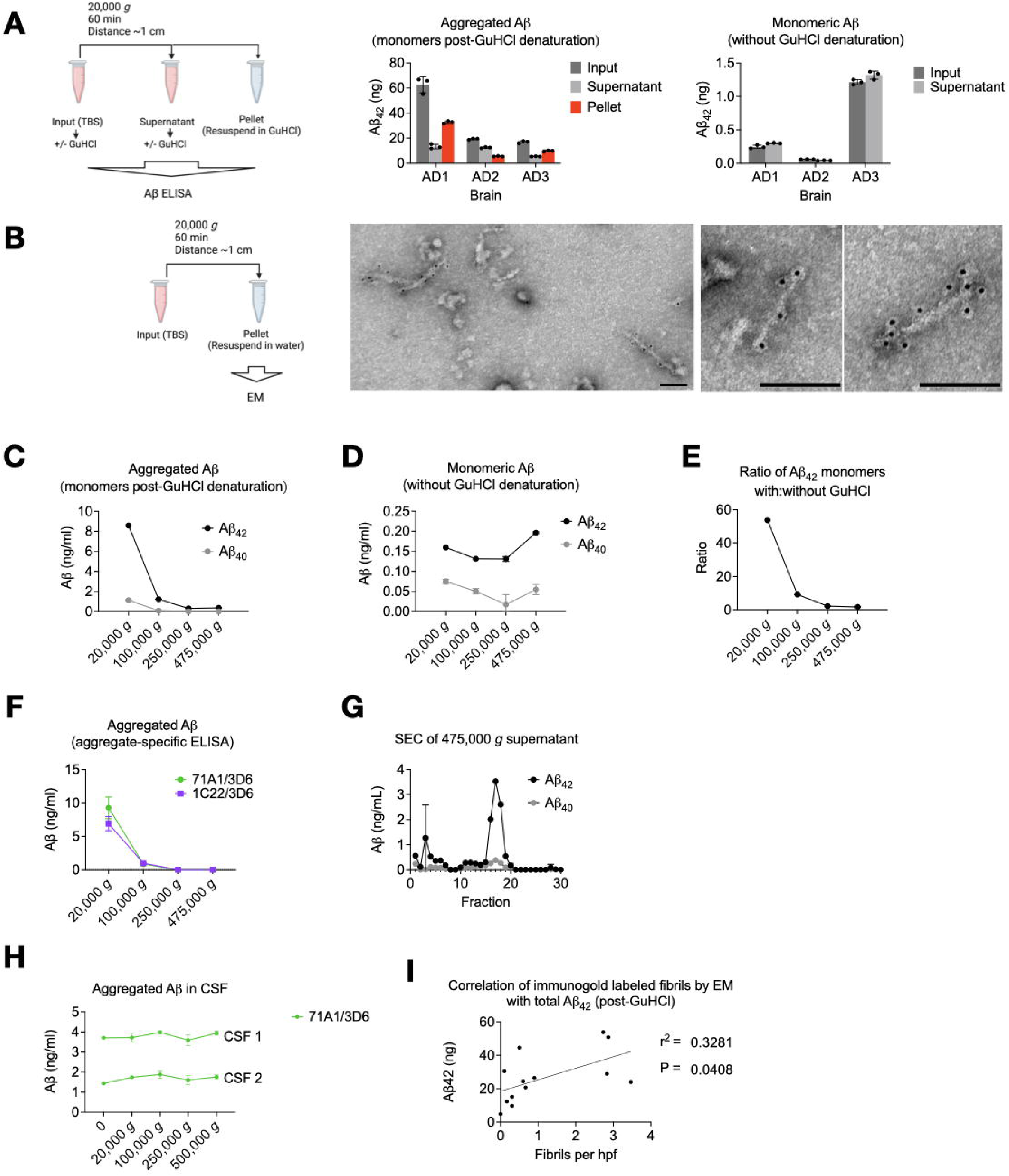
Aqueous soaking extracts of AD brain contain insoluble and fibrillar. **A**β. **(A)** Aliquots of aqueous soaking extracts from ultracentrifugal supernatants of AD cortex (see Methods and Table 1) were thawed and spun at 20,000 *g* in a tabletop centrifuge, and Aβ was quantified by ELISA with (center) and without (right) 5 M GuHCl. Note the different y-axis scales. GuHCl is a chaotropic agent which denatures Aβ aggregates and allows detection by a monomer-preferring ELISA. Aβ detectable without GuHCl denaturation is monomeric. ELISAs revealed that aggregated Aβ, (center) but not Aβ monomers (right), could be re-pelleted. **(B)** EM of fibrils in the pellets of re-centrifuged aqueous soaking extracts of brain AD7 were labeled by anti-Aβ N-terminal antibody D54D2. Scale bar = 100 nm. **(C-F)** Ultracentrifugation pellets high MW Aβ aggregates. A TBS soaking extract of brain AD6 was prepared by mincing cortex, soaking in TBS for 30 min, followed by a 10 min spin at 2,000 *g* (Table 1, steps 1-4). This supernatant was then divided and ultracentrifuged in a TLA100.3 rotor (pelleting distance ~2 cm) at indicated *g* forces, and Aβ quantified in the supernatant. Monomer-preferring ELISA performed with (C) or without (D) denaturation with GuHCl showed that Aβ aggregates but not native monomers were depleted as the *g*-force increased, as did the ratio of Aβ concentration with:without GuHCl (E). Two different aggregate-preferring ELISAs showed abolished signal as the *g*-force increased. Superdex 200 Increase size exclusion chromatography of the 475,000 *g* supernatant revealed a low MW Aβ profile. **(H)** In contrast to brain, CSF Aβ aggregates were not depleted as *g*-force increased. **(I)** The amount of Aβ measured in soaking extracts after GuHCl denaturation correlated with the number of fibrils present in the pellet after centrifugation and manual blinded counting by immunogold EM. Line represents simple linear regression. Drawings made with Biorender.com.

Aβ fibrils were not contaminants from the ultracentrifugation pellets because they were also present when only the top ~20% of the supernatants was collected (Fig S2A). To rule out that the sedimentable fibrils we observed had adhered to the surface of the minced tissue (*e*.*g*., on the cut surfaces of plaques) rather than diffused out of the tissue into the soaking buffer, we rinsed the minced brain bits with 15 volumes of extraction buffer through a 100 μm filter prior to soaking. We found no reduction in the amount of extracted Aβ aggregates measured biochemically (Fig S2B) or as amyloid fibrils seen by EM. Further arguing in favor of diffusion of fibrils from the minced brain bits into the aqueous buffer rather than carryover from the cut tissue surface was the observation (also documented in Fig S2 of Hong *et al*.^21^) that diffusion of Aβ into the buffer required 15-30 min to plateau, a similar rate as the diffusion of control soluble brain proteins.

### Sufficient centrifugal force can pellet HMW Aβ aggregates

When accounting for the pelleting distance, our initial and re-centrifugation protocols were approximately equal in pelleting efficiency: the pelleting distance in thin-wall tubes in the SW41Ti (initial soaking extract) is ~9 cm when full (Table 1, Step #5), whereas it is only ~1 cm for 1 ml in a microcentrifuge tube at a 45-degree angle in the benchtop centrifuge (Table 1, Step #7). Thus, the benchtop centrifugation should accomplish the same pelleting efficiency at an ~9-fold lower *g-*force. It was therefore possible that if higher *g-*forces had been used initially, the fibrils would not have been recoverable from the supernatant during re-centrifugation. We modified the initial ultracentrifugation step, using the fixed-angle TLA100.3 rotor (pelleting distance ~2 cm) at increasing *g*-forces, instead of the SW41Ti (pelleting distance ~9 cm). We found that *g*-forces ≥250,000 *g* across the shorter distance depleted nearly all biochemically detectable aggregated Aβ from the supernatants (Fig 1C) but retained native monomers (Fig 1D). In 475,000 *g* supernatants, addition of GuHCl denaturation released 1.8-fold more Aβ monomers than without GuHCl, whereas in 20,000 *g* supernatants, GuHCl addition released 54-fold more Aβ monomers (Fig 1E). This suggests that most Aβ in the 475,000 *g* supernatants consisted of soluble native monomers and covalent dimers.^3,20^ To verify that Aβ aggregates were quantitatively depleted from supernatants obtained at or above 250,000 *g*, we used high-sensitivity sandwich ELISAs with two different published aggregate-preferring capture antibodies^24,25^ that do not require GuHCl denaturation (Fig 1F). Neither assay could detect aggregates in the ≥250,000 *g* supernatants. Moreover, residual Aβ in the 475,000 *g* supernatants exhibited a LMW SEC profile (Fig 1G).

We conclude that most Aβ, and all its HMW aggregates from aqueous brain extracts are made of particles in suspension, consistent with at least some of them being fibrils. We then compared this finding to Aβ in cerebrospinal fluid. Detection of CSF Aβ by our aggregate-specific ELISA was unaltered regardless of the centrifugal force used (Fig 1H), suggesting that the Aβ aggregates detected in human CSF were soluble, in contrast to those in aqueous extracts of AD brain.

### Aβ fibrils in aqueous soaking extracts do not appear to form during sample preparation

We next sought evidence of *postmortem* artifact: that the fibrils were not present during life but polymerized from smaller non-fibrillar aggregates during the *postmortem* interval or during sample processing. We continued using the standard soaking extract protocol using the SW41Ti rotor at 200,000 *g*,^21^ followed by re-centrifugation in the tabletop centrifuge at 20,000 *g* (Table 1).^21^

One possibility was that Aβ polymerization occurred along the plastic tube surface during re-centrifugation of soaking extracts. We performed re-centrifugation in which standard soaking extracts were spun and their supernatants moved to new tubes at intervals. The Aβ in each pellet was quantified by ELISA following GuHCl denaturation. If nucleation of new fibrils had occurred, we would have expected a lag in the accumulation of Aβ in the pellets, whereas pelleting of pre-existing particulates would be linearly time-dependent until they had been depleted from the supernatants. We observed no lag phase, suggesting that pelleted Aβ did not arise from nucleation of new fibrils along the plastic tubes (Fig S2C, D). Storage of standard soaking extracts in siliconized or BSA-coated tubes also had no effect compared to standard polypropylene (Fig S2E).

The next possibility was that Aβ polymerization occurred during freezing and thawing of the soaking extracts. When proceeding directly from the standard SW41Ti (Table 1, step 5) to a second spin in the benchtop centrifuge (step 7) without freezing, Aβ aggregates as measured by ELISA could still be pelleted (Fig S3A), and fibrils were observed by immuno-EM (Fig S3B). Nevertheless, we also found that freezing a freshly prepared extract at −80°C for storage, as typically performed, resulted in re-pelleting more Aβ from a soaking extract once thawed compared to storage at 4°C (Fig S3C). We also found that the re-pelleting of Aβ was concentration-dependent: when the SW41Ti ultracentrifugation supernatants were diluted at least four-fold before freezing, less aggregated Aβ could be re-pelleted after thawing (Fig S3D). To assess further whether polymerization occurred during the freeze/thaw cycle, we added inhibitors of Aβ aggregation, including aggregate-binding antibodies, 0.45% CHAPS,^26^ and *scyllo*-inositol,^27,28^ immediately after the initial ultracentrifugation of the soaking extracts and before freezing the supernatants. None of these agents reduced the amount of Aβ that could be re-pelleted after thawing (Fig S3E, F). We also expected that if fibrils had formed *ex vivo* from monomers and/or soluble LMW aggregates during freeze/thaw, then filtering the soaking extracts immediately after the original ultracentrifugation and before freezing should have had no effect on the recovery and pelleting of Aβ. We found, however, that freshly prepared soaking extracts were depleted of Aβ aggregates by a 20-nm pore size alumina filter (Fig S3G) but passed through a 100-nm filter to control for non-size dependent binding to the filter. These results implied that most Aβ assemblies were at least 20 nm in size prior to freezing the original soaking extract. SEC of a freshly prepared TBS extract immediately after the original ultracentrifugation (without freezing) revealed that Aβ still eluted in the void volume of a Superdex 200 Increase column (~≥500 kDa) (Fig S3H). Together, these results suggest that Aβ aggregates were HMW and likely fibrillar prior to freezing and storage of the original soaking extracts. Increased pelleting of Aβ fibrils after freeze/thaw may be due to increased clumping of existing fibrils but probably not to formation of new fibrils.

Another possibility was that polymerization occurred after death but before tissue freezing. To exclude this artifact as best as possible, we prepared a soaking extract and applied it to the Superdex 200 Increase column within 8 h of an AD patient’s death without freezing either the tissue or the extract. Again, virtually all the ELISA-detectable aggregated Aβ eluted in the void volume of a Superdex 200 Increase column (Fig S3I-K), and fibrils could be pelleted during re-centrifugation (Fig S3L).

### Fibrils account for a substantial proportion of high-molecular weight Aβ aggregates in soaking extracts

We manually counted the D54D2-immunoreactive fibrils in the re-centrifugation pellets from 13 AD brains, using 30 high-powered EM fields each. Using linear regression, we found a correlation between the average number of fibrils per field and the Aβ levels measured by ELISA after GuHCl denaturation in those same extracts (Fig 1I).

### Fibrils from soaking extracts of AD brain have the same cryo-EM structures as those from sarkosyl-insoluble preparations

The cryo-EM structures of Aβ fibrils from the sarkosyl-insoluble fractions of AD brains were recently determined.^16^ We now asked if the fibrils we identified in aqueous soaking extracts resembled those from sarkosyl-insoluble homogenates. Compared to Aβ fibrils from sarkosyl-insoluble fractions, the soaking extract-derived pellets showed shorter fibrils that were less clumped together, compatible with their relative resistance to pelleting. Cryo-EM structure determination of soaking extract-derived fibrils from two cases of sporadic AD (AD7 and AD11) revealed identical structures to those reported previously for Aβ fibrils from sarkosyl-insoluble preparations.^24^ (Fig 2). Case AD7 had only Type I fibrils, whereas case AD11 had both Type I and Type II fibrils, as defined previously.^16^

**Fig 2.**
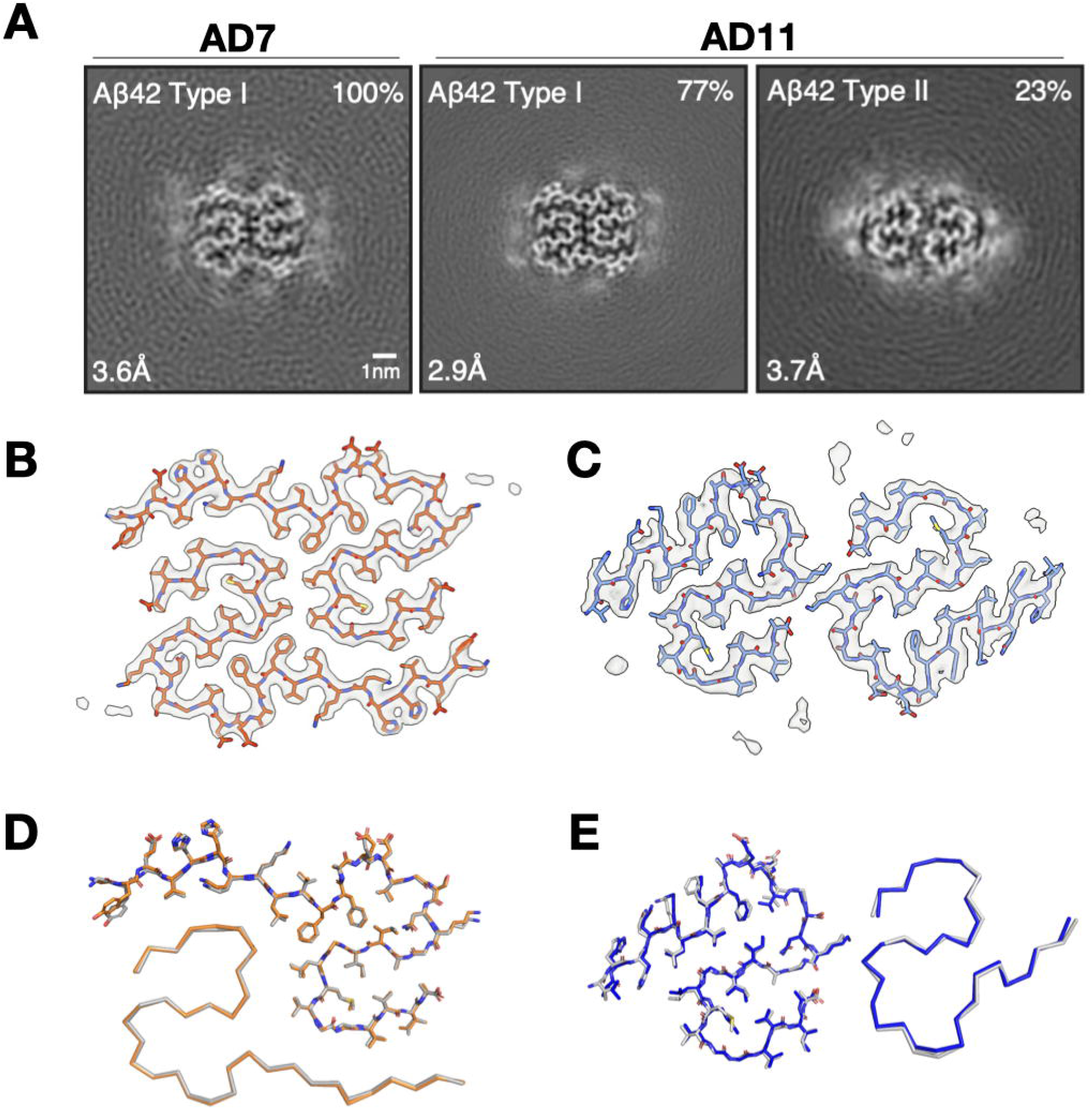
Aβ fibrils from aqueous extracts of AD brains have the same cryo-EM structures as amyloid fibrils from sarkosyl-insoluble homogenates. **(A)** XY-cross-sections of cryo-EM maps of Aβ filaments from soaking extracts of cases AD7 and AD11. For each map, a sum of the reconstructed densities for several XY-slices, approximating one β-rung, is shown. Aβ filament types I and II^16^ are indicated at top left; the percentages of Type I and Type II filaments relative to the total of imaged filaments are shown at top right. Resolutions at bottom left. Scale bar = 1 nm. **(B, C)** Cryo-EM density maps (in transparent grey) and atomic models for Aβ Type I (in orange in B), Aβ Type II (in blue in C). **(D, E)**, Comparison of the cryo-EM structures of Aβ Type I (B) filaments and Aβ Type II (C) filaments extracted from aqueous soaking extracts vs. sarkosyl-insoluble homogenates (in grey for Type I and Type II filaments) from brains of AD patients.^16^ Structures are shown as sticks (above) for one protofilament and as ribbons (below) for the other.

Because they were present in the same re-centrifuged soaking extracts, we also determined the cryo-EM structures of the tau filaments. They were identical to those reported previously for PHFs from sarkosyl-insoluble fractions of AD brains (Fig S4).^29^

### Lecanemab labels Aβ fibrils from soaking extracts

Lecanemab was derived from monoclonal antibody 158 that was selected to bind synthetic aggregates of Aβ bearing the AD-causing *APP* “Arctic” mutation E22G.^1,15^ Synthetic Aβ_40_[E22G] forms more soluble aggregates than the wild-type Aβ_40_ peptide *in vitro*, and the mechanism of action of lecanemab is thought to relate to its preference for small, soluble Aβ aggregates over amyloid fibrils.^30^ Lecanemab has been shown to immunoprecipitate Aβ from aqueous AD brain extracts,^15^ so we asked whether lecanemab can bind the fibrils we observed. We found lecanemab could decorate Aβ fibrils (Fig 3A, B) that had been re-pelleted from AD soaking extracts. We also examined antibody h1C22, which we had previously reported to bind better to synthetic Aβ aggregates than to monomers and fibrils,^7,23,24^ and we again observed labeling of Aβ fibrils (Fig 3C-E).

**Fig 3.**
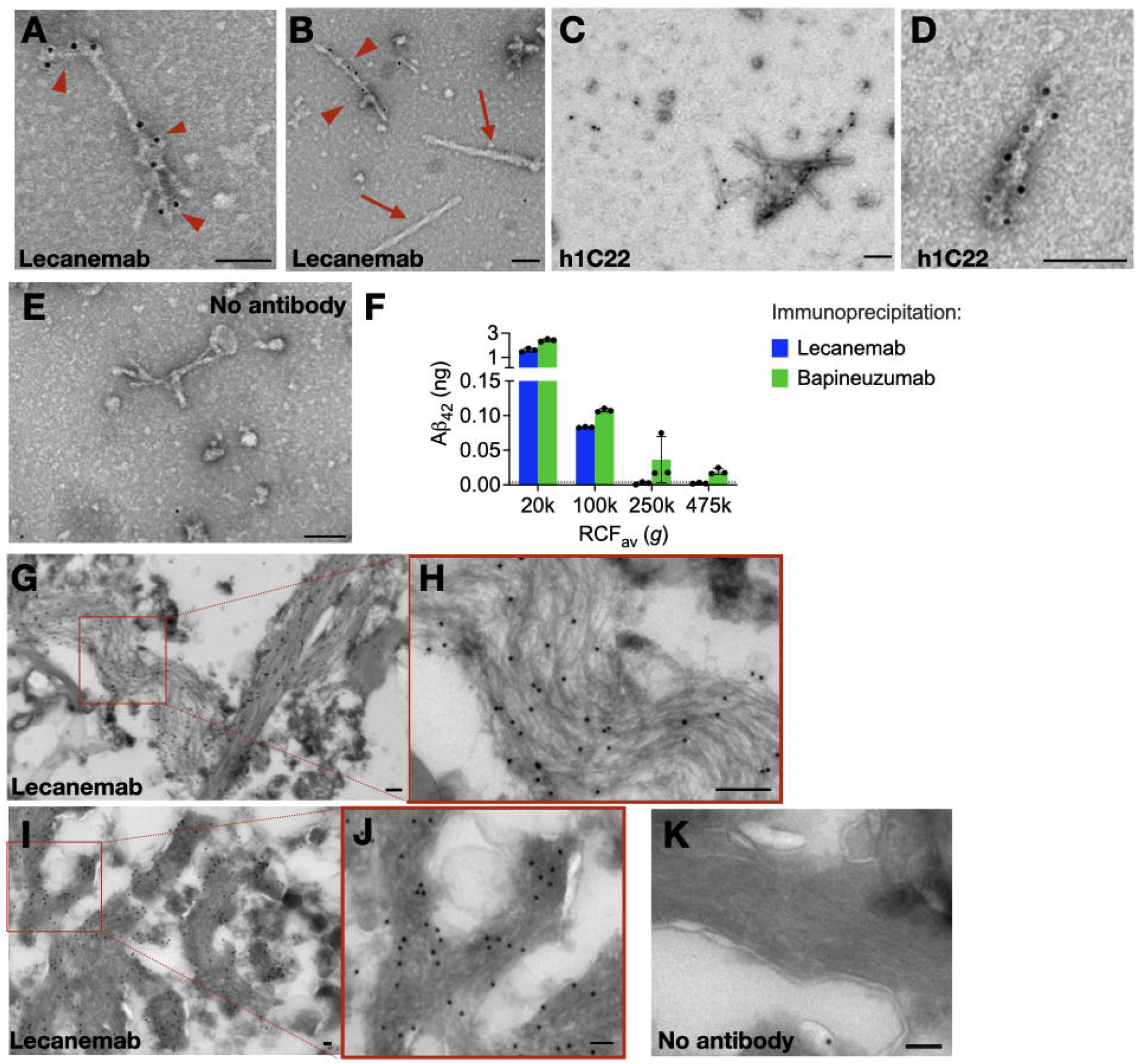
Lecanemab decorates Aβ fibrils from Alzheimer’s disease brains. **(A,B)** Immunogold labeling of pelleted material from the re-centrifugation of AD7 TBS soaking extract reveals immunoreactivity of Aβ fibrils with lecanemab (arrowheads). PHF in the pellet did not react with lecanemab (arrows). **(C, D)** Immunogold labeling with h1C22 Aβ aggregate-preferring antibody shows similar decoration of soaking extract fibrils. **(E)** Omitting primary antibody shows no labeling of fibrils, but infrequent background protein A-gold particles. **(F)** ELISA quantification of Aβ_42_ immunoprecipitated from AD6 aqueous extracts followed by washing, elution and denaturation in 5 M GuHCl. Bapineuzumab, but not lecanemab, still immunoprecipitated Aβ from the 250,000 *g* and 475,000 *g* (TLA100.3, pelleting distance ~2 cm) aqueous soaking extract supernatants. **(G-I)** Immunogold labeling of paraformaldehyde-fixed ultrathin cryosections from AD11 (G,H) and AD7 (I,J) brains reveals labeling of amyloid fibrils by lecanemab *in situ*. (**K**) Protein A gold alone without lecanemab (brain AD7) shows minimal background labeling. Scale bar = 100 nm.

Since lecanemab labeled Aβ fibrils from aqueous brain extracts, we hypothesized that it would only immunoprecipitate Aβ from extracts prepared at *g*-forces that do not deplete aggregates detectable by other means, such as aggregate-preferring ELISAs that capture Aβ with 1C22 or 71A1 (Fig 1F).^25^ We found that lecanemab could only immunoprecipitate Aβ from aqueous AD brain supernatants that had been ultracentrifuged at ≤100,000 *g* in the TLA100.3 rotor (pelleting distance ~2 cm) (Fig 3F), similar to 1C22 (Fig. 1F). In contrast, bapineuzumab, which binds both monomeric and aggregated Aβ, could still immunoprecipitate some Aβ from 250,000 and 475,000 *g* supernatants.

Lecanemab has been shown to label amyloid plaques by immunohistochemistry at the light microscopic level.^31^ To determine if it stains plaque fibrils, we prepared ultrathin cryosections from AD cortex using minimal fixation, so as to preserve antibody epitopes. Although the cytoarchitecture was not well preserved as expected, we observed lecanemab labeling of amyloid fibrils *in situ* (Fig 3G-J) and minimal background from protein A gold alone (Fig 3K).

## Discussion

Characterization of Aβ aggregates in aqueous extracts of AD brain has been difficult, but our results suggest that at least some are identical in tertiary structure to insoluble amyloid fibrils. After gentle soaking extraction (diffusion) from frozen or fresh AD cerebral cortex in aqueous buffer, almost all the Aβ aggregates were retained in the void volume of a Superdex 200 Increase column, agreeing with previous studies.^18,19,21^ The Aβ aggregates in these soaking extracts could be re-pelleted, and the resultant pellets contained amyloid fibrils that were labeled by several anti-Aβ antibodies, including the putatively soluble aggregate-preferring lecanemab.^15,32^ The aqueously extracted fibrils had the same cryo-EM structures as those of Aβ fibrils from the sarkosyl-insoluble pellets of homogenized AD cortex.^16^ While we cannot rule out the presence of some non-fibrillar HMW Aβ assemblies, a correlation between total Aβ levels measured by ELISA and the number of Aβ fibrils seen by EM (Fig. 1I) suggests that amyloid fibrils account for a substantial proportion of HMW aggregates that diffuse into aqueous extracts.

An unavoidable limitation of this work is the need to examine AD brains *postmortem*. Although we attempted to mitigate the likelihood of artifacts, we cannot completely exclude disassembly of some plaques into free-floating fibrils after death that then entered our soaking extracts. We also cannot completely exclude the *ex vivo* aggregation of monomers or LMW aggregates (*e*.*g*., dimers) into fibrils. While we addressed this possibility by testing the effects of small-pore filters and inhibitors of Aβ aggregation, and by using fresh tissue when possible, some *ex vivo* aggregation may have occurred.

If small Aβ fibrils diffuse through human brain, they may be in equilibrium with amyloid plaques. By immunohistochemistry, APP transgenic mice show halos surrounding methoxy-XO4-labeled amyloid plaques that are immunoreactive with Aβ aggregate-preferring antibodies.^33^ The halos may consist of Aβ fibrils too loosely packed to shift the fluorescence spectra of such dyes and having more access to local synaptic endings than do the fibrils within compacted plaque cores. Supporting this model is our observation that aqueously diffusible fibrils have the same cryo-EM structures as detergent-insoluble Aβ fibrils. Experiments in mice in which the microglial AD risk gene *TREM2* was knocked out, or else microglia were depleted, have shown that compaction of Aβ by microglia may prevent fibril access to dendrites and synapses.^34–36^ Soaking brain extracts prepared identically to those studied here (Table 1) have been shown to confer neurotoxicity: impaired synaptic plasticity in mouse hippocampal slices and neuritic dystrophy (reduced neurite length and branching) in cultured human neurons.^21^ Moreover, we previously found that HMW aggregates from AD brain had less bioactivity toward both synapses and microglia than did the LMW aggregates (including dimers) into which they disassembled.^19^ We now propose that many of these HMW assemblies are short fibrils; thus, partial disassembly of diffusible Aβ fibrils may be required to exert neurotoxicity.

Our results have implications for Aβ-targeting therapies. Lecanemab was derived from mouse monoclonal antibody 158, which is reported to preferentially recognize intermediate and lower molecular weight forms of synthetic Aβ aggregates.^15^ However, lecanemab also reduces amyloid PET signal, a marker of fibrillar Aβ,^37^ and labels plaques by immunohistochemistry. ^31^ Based on our finding that aqueous AD brain extracts contain Aβ fibrils that are decorated by lecanemab, we speculate that lecanemab’s binding target in the human brain may include diffusible fibrils with the same structure as those in insoluble plaques. It remains possible that lecanemab’s target may also include non-fibrillar aggregates because we could not exclude their presence.

Finally, while we focused on Aβ, we also observed PHF tau present in our soaking extracts. Others have observed that tau seeding activity in aqueous extracts correlates with molecular weight by SEC and inversely with the centrifugal force generating the extracts, implying that larger and heavier tau aggregates are more seed-competent.^12,13^ Based on our cryo-EM analysis, we suspect that these preparations contained short tau fibrils with the same fold as PHFs from sarkosyl-insoluble preparations.^38^ Supporting this model are the observations that recombinant tau assemblies shortened by sonication were not pelleted by ultracentrifugation,^39^ and that tau fibrils were the most seed-competent in brain extracts from transgenic mice overexpressing human mutant P301S tau.^40^ Future efforts aimed at the rational design of therapeutic agents that bind and clear tau seeds should consider that they may be PHF.

## Acknowledgements

We thank Drs. Matthew Frosch, Mel Feany and Michael B. Miller for assistance in brain tissue allocation. We thank Alexandra Golby, MD, and Mary-Beth Anketell, NP, MSN, for help in acquiring patient CSF. Funded by NIH K08NS128329 (AMS), P01AG015379 (DJS), and RF1AG006173 (DJS); Alzheimer’s Association AACSF-21-849687 (AMS); the Davis Alzheimer’s Prevention Program (DJS); and Medical Research Council MC_UP_A025_1013 (SHWS) and MC_U105184291 (MG). DJS is a director of Prothena Biosciences; the other authors declare no competing interests.

## Figure Legends

**Fig S1.**
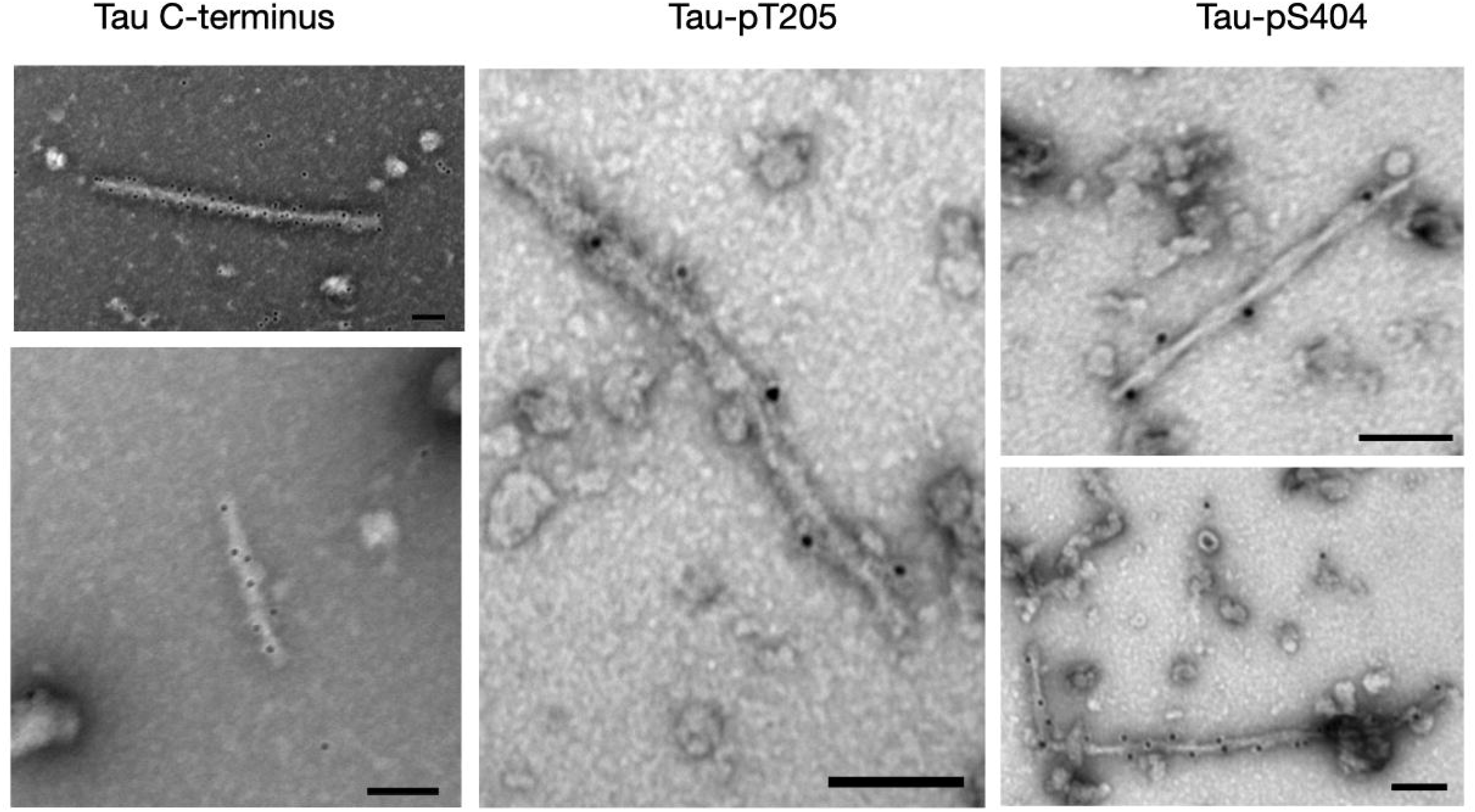
Presence of tau filaments in aqueous AD brain extracts. Immunogold labeling of pellets from the 20,000-*g* tabletop re-centrifugation of TBS soaking extracts of brains AD12 and AD8 revealed PHFs immunoreactive with antibodies to the tau C-terminus and to tau phosphorylated at T205 or S404. Scale bar = 100 nm.

**Fig S2.**
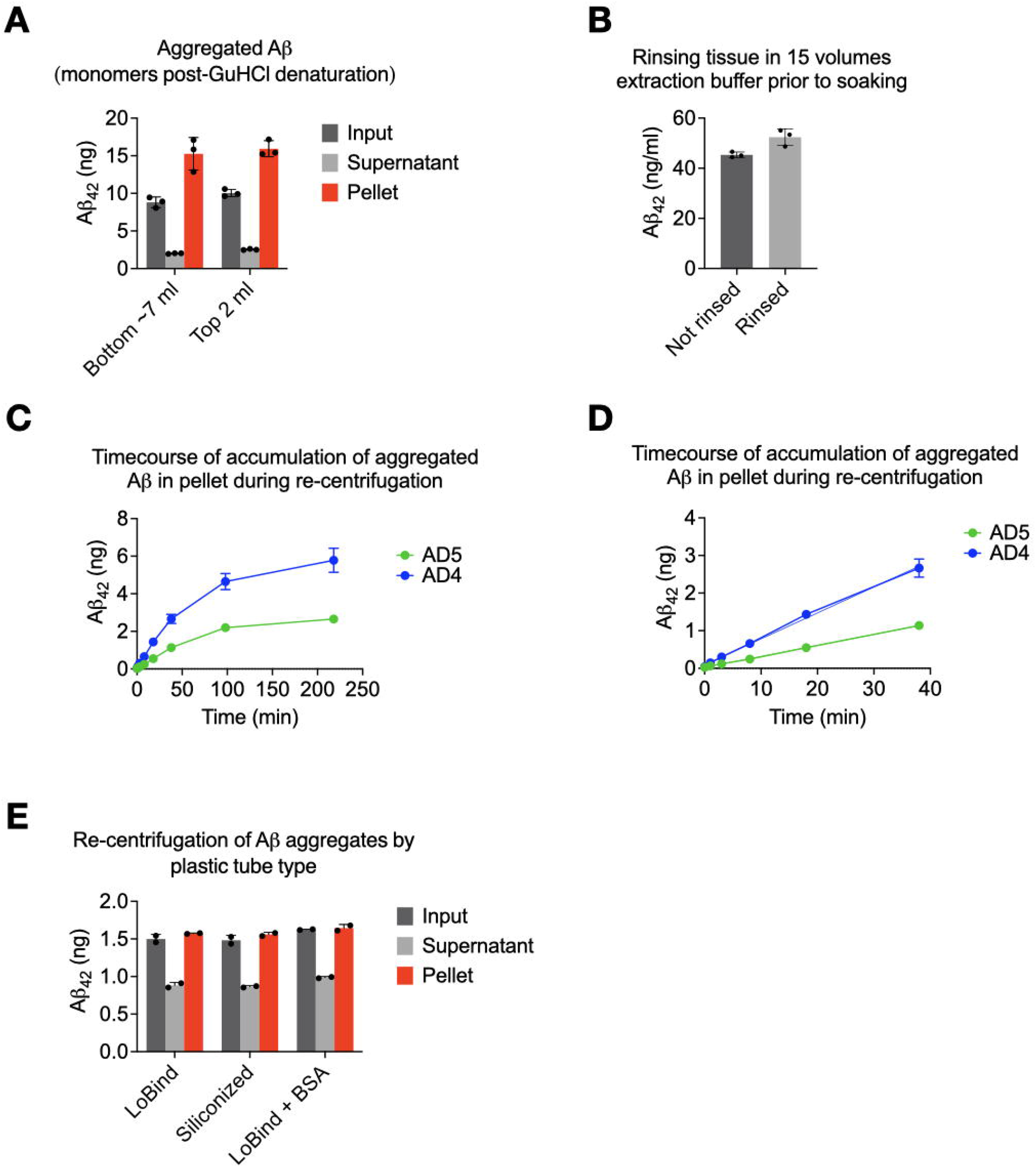
Aβ fibrils in soaking extracts are not contaminants from the ultracentrifugation pellet or adhered to the tissue, and do not arise due to plastic-induced polymerization. **(A)** Equivalent quantities of Aβ aggregates could be pelleted from the top 2 ml and bottom 7 ml portions of the ultracentrifugal supernatant of brain AD6. **(B)** Rinsing minced brain bits of AD12 in 15 vol of aqueous extraction buffer to remove adherent Aβ aggregates prior to soaking does not decrease Aβ recovery. **(C)** Centrifugation of two thawed AD TBS soaking extracts reveals the absence of a lag phase in the accumulation of Aβ in the pellet. The soaking extracts were centrifuged sequentially for progressively longer times, with each pellet quantified for Aβ_42_, to search for a lag phase that could indicate nucleation. No lag phase was observed. **(D)** Expanded view of the first few timepoints of (C) confirms linear pelleting without a lag phase. Storage of frozen AD4 TBS soaking extracts in siliconized or BSA-coated tubes does not alter amount of Aβ recovered in the pellets vs. standard polypropylene Protein LoBind tubes.

**Fig S3.**
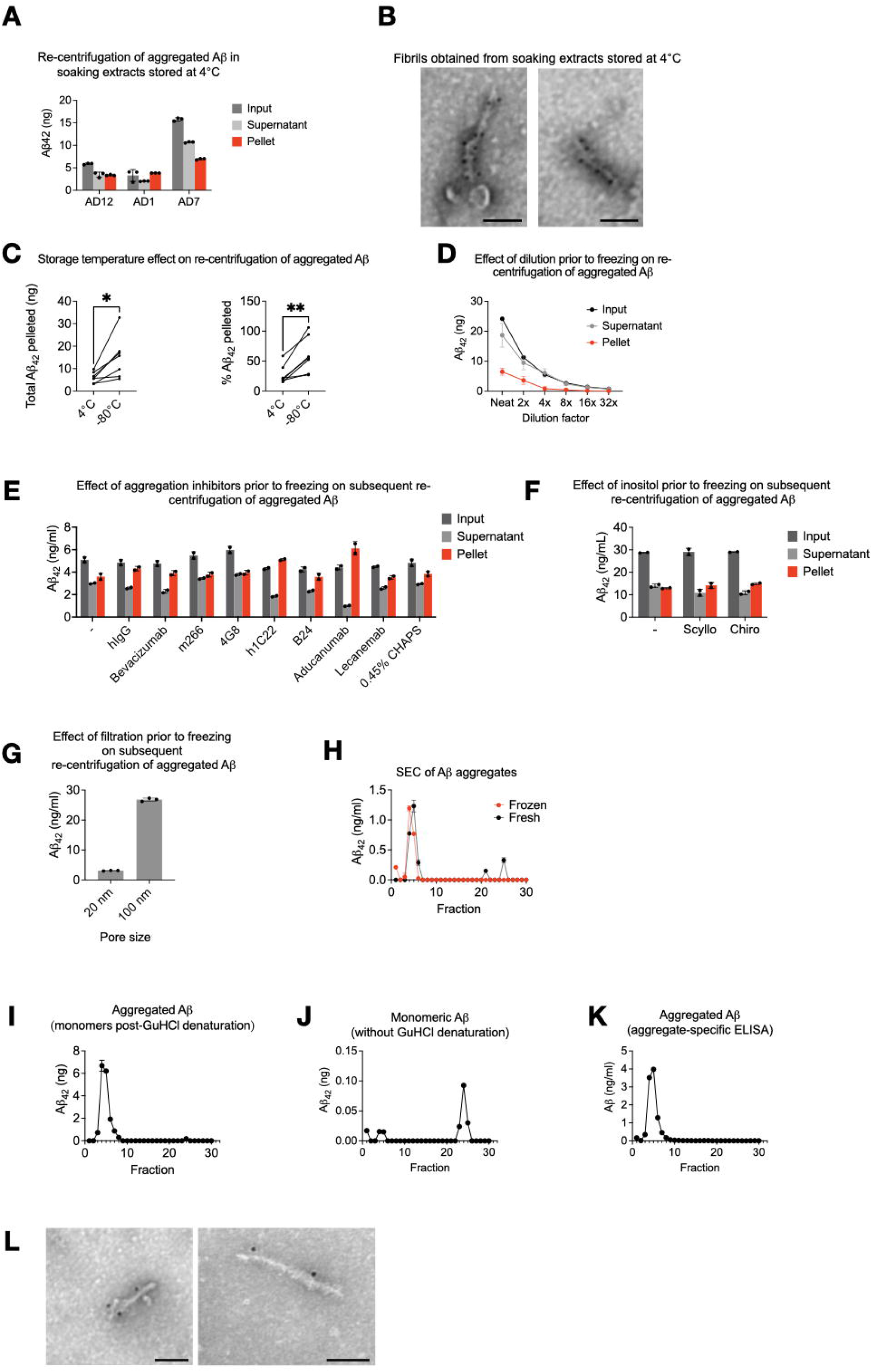
No evidence that Aβ fibrils from soaking extracts form *postmortem*, but the amount recovered is enhanced by freeze-thaw. **(A)** The SW41Ti supernatant from three different AD brain soaking extracts was immediately aliquoted and spun in a tabletop centrifuge at 20,000 *g without freezing*. Aβ_42_ ELISA reveals the ability to pellet Aβ (red). (**B**) Immunogold EM of pellet from the AD7 soaking extract in (A) (no freezing) reveals Aβ-immunoreactive (D54D2) fibrils. **(C)** More Aβ (by ELISA post-GuHCl) is re-pelleted from soaking extracts frozen at −80°C than kept at 4°C (N=7 AD brains). Left, total Aβ pelleted; right, Aβ pelleted as % of Aβ input. *P<0.05, **P<0.01 by two-tailed paired student’s t-test. **(D)** Serial dilution of original TBS soaking extract (AD7) immediately after removal from the ultracentrifuge and before freezing diminishes recovery of Aβ_42_ upon re-pelleting. **(E)** Addition of 7 different Aβ monoclonal antibodies (10 µg/ml) or CHAPS at 0.45% to TBS soaking extract (AD4) immediately after the ultracentrifugation step did not decrease subsequent re-pelleting after freeze-thaw storage. **(F)** Addition (2 mM) of anti-Aβ *agg*regation agent *scyllo*-inositol (or its inactive stereoisomer *chiro*-inositol) to TBS soaking extract (AD9) immediately after the ultracentrifugation did not decrease subsequent re-pelleting after freeze-thaw storage. **(G)** A 20-nm alumina (Whatman Anotop) filter removed more Aβ aggregates than a 100-nm filter (control) before freezing a freshly prepared TBS soaking extract of brain AD8, suggesting that most Aβ particles were larger than 20 nm before freezing. **(H)** Aβ mostly elutes in the void volume of a Superdex 200 Increase column both before and after freezing (−80°C) freshly prepared TBS soaking extract of brain AD4. **(I-K)** A soaking extract from brain AD10 was made and analyzed by size exclusion chromatography (Superdex 200 Increase) by <8 h after death. Monomer-preferring ELISA performed with (I) and without (J) denaturation with GuHCl shows Aβ_42_ aggregates mostly in the void volume, as confirmed with an Aβ aggregate-preferring ELISA (K). **(L)** After freezing soaking extracts and later thawing and re-pelleting, D54D2-immunoreactive fibrils were present in the pellets. Scale bar = 100 nm.

**Fig S4.**
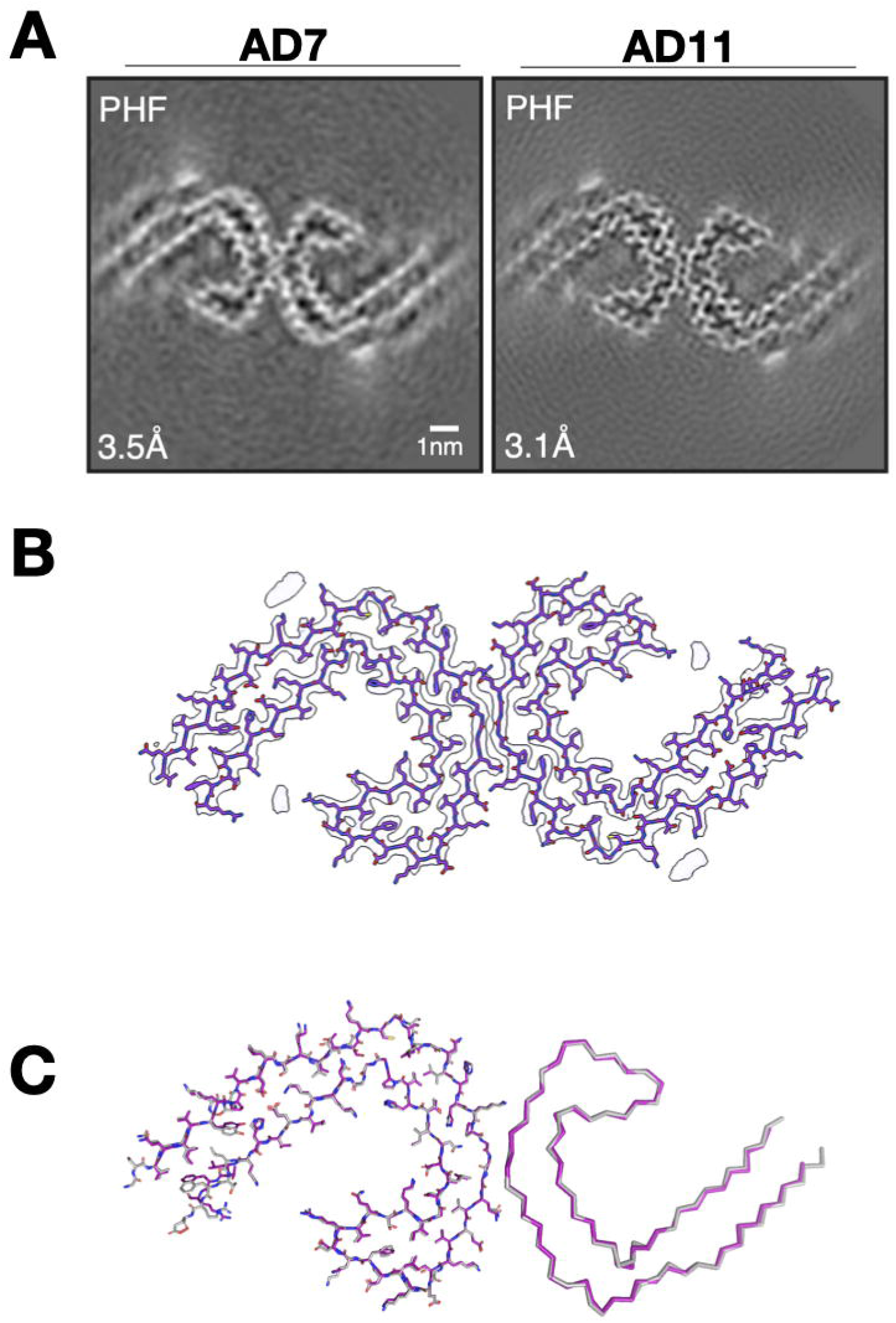
Tau fibrils from soaking extracts of AD cortex have the same structures as those of PHFs in sarkosyl-insoluble homogenates. **(A)** Cryo-EM maps of PHFs from soaking extracts of AD7 and AD11. For each map, a sum of the reconstructed densities for several XY-slices, approximating one β-rung, is shown. Resolutions at bottom left. Scale bar = 1 nm. **(B)** Cryo-EM density map (in grey) and atomic model for PHF (in purple). **(C)** Comparison of the cryo-EM structures of PHFs extracted from soaking extracts (in magenta) and from sarkosyl-insoluble homogenates (in grey) of AD brains.^29^ Structures are shown as sticks for one protofilament (above) and as ribbons for the other protofilament (below).

## Materials and Methods

### Aqueous soaking extraction of Aβ and tau fibrils from Alzheimer’s disease brains and their re-centrifugation

The soaking method for aqueous extraction of diffusible Aβ aggregates from AD brain has been described previously^21,22^ and is depicted in Table 1. Brain donor demographic and pathologic information is described in Table S1. Grey matter was dissected from fresh or frozen cortical regions of individuals with AD and minced using a McIlwain tissue chopper (razor blade) set at 0.5 mm. The resultant brain bits were soaked for 30 min in a 1:5 ratio (w:v) of TBS extraction buffer (25 mM Tris supplemented with 150 mM NaCl, 5 μg/ml leupeptin, 5 μg/ml aprotinin, 2 μg/ml pepstatin, 120 μg/ml 4-benzenesulfonyl fluoride hydrochloride, 5 mM NaF, pH 7.2) in 50-ml Eppendorf Protein LoBind tubes, and then spun at 2,000 *g* in a Fiberlite F14-14 × 50cy rotor in a Sorvall Lynx 6000 centrifuge for 10 min at 4°C (pelleting distance ~4 cm depending on volume). The top ~90% of the supernatants were transferred to thin-wall polypropylene tubes (Beckman catalog #331372) and spun using an SW41Ti rotor at 40,000 rpm for 110 minutes in an Optima L90K ultracentrifuge at 4°C (pelleting distance ~9 cm). The top ~90% supernatants were retained and frozen at −80°C in 1 ml aliquots. For re-centrifugation (see Results), the frozen aliquots were thawed and spun in an Eppendorf 5417R centrifuge equipped with an F-45-30-11 rotor or else a TOMY MX-307 centrifuge equipped with an AR015-24 rotor at 20,000 *g* for 2 h at 4°C (pelleting distance ~1 cm). Prior to this re-centrifugation, a 50 μl portion of each aliquot was retained for Aβ ELISA. The final supernatants, as well as the associated pellets, were collected and retained for ELISAs and electron microscopy (EM). For Aβ42 monomer ELISAs, pellets were resuspended and denatured in 50 μl TBS containing 5 M GuHCl and 5 mM EDTA. Similarly, for Aβ42 monomer ELISAs on the original soaking extracts (inputs) and their recentrifuged supernatants, each was denatured by mixing them at a 2:5 ratio (v:v) with TBS containing 7 M GuHCl and 5 mM EDTA, to reach a final GuHCl concentration of 5 M. For analysis by negative-staining EM, final pellets were resuspended in 15 μl cold water.

### ELISAs

Aβ monomer-preferring ELISAs were home-brew immunoassays run on the Meso Scale Discovery (MSD) platform, as described.^23^ Samples containing GuHCl were diluted before ELISA, such that the GuHCl concentration was <0.25 M, to avoid interference with the antibody reaction. All steps were at room temperature. Plates were coated with capture antibody in PBS overnight and blocked in 5% MSD Blocker A in TBS with 0.05% Tween-20 (TBS-T) for 1 h. Samples were then applied for 1.5 h after being diluted in 1% Blocker A in TBS-T. Plates were washed 3 times in TBS-T, then biotinylated detector antibody and MSD Strepavidin-Sulfotag (1:5000) were applied for 1.5 h. After three more washes in TBS-T, the plates were detected with 2x MSD read buffer. The lower limit of quantification (LLoQ) was defined as the lowest standard with a luminescence value at least twice the blank average, and the lower limit of detection (LLoD) was defined as the lowest standard with values greater than the blank average plus twice the blank standard deviation. Aβ aggregate-preferring ELISAs were home-brew assays run on the Millipore SMCxPRO platform, as described previously.^25^ Antibodies, sources, and concentrations are listed in Table S2.

### Negative staining and immunoelectron microscopy

Carbon-coated grids (EMS) were glow-discharged at 25 mA for 20 s. For liquid samples (resuspended final pellets from soaking extracts), 5 μl were adsorbed to the grid for 1-5 min. Excess liquid was blotted off with filter paper (Whatman #1). Grids were then floated on a drop of water, blotted again, and stained with 1% uranyl acetate for 15 s. Excess stain was blotted off with filter paper. For immunostaining after their adsorption to the grids, samples were blocked in 1% BSA for 10 min. Antibodies diluted in 1% BSA were then added for 20 min and washed 3 times in PBS. Protein A-gold (Cell Biology UMC Utrecht) in 1% BSA was applied for 30 min, followed by two washes in PBS and four washes in water. All grids were examined on a JEOL 1200EX transmission electron microscope equipped with an AMT 2k CCD camera. Except for lecanemab, primary antibodies were purchased from Cell Signaling Technologies and diluted as follows: Anti-Aβ N-terminus (D54D2) 1:10; anti-tau C-terminus (D1M9X) 1:20; anti-tau pS404 (D2Z4G) 1:50; anti-tau pT205 (E7D3E) 1:50. Lecanemab was obtained from Sanofi and used at a final concentration of 13.3 µg/ml. To quantify fibrils, thirteen 0.9-ml soaking extracts were centrifuged for 2h at 20,000 *g* in an Eppendorf FA-45-30-11 rotor, and the pellets were resuspended in 20 μl water. Samples were immunolabeled with D54D2 as above, and 30 micrographs were taken at random by one experimenter (AMS) from well-stained portions of each grid at 25,000x magnification. The image order was randomized by a second experimenter (ALM) and given to a third blinded experimenter (ME) who counted the number of immunolabeled fibrils.

For ultrathin cryosections, AD brain tissues were fixed (fresh or after freezing) in 4% paraformaldehyde and stored at 4°C. Prior to freezing in liquid nitrogen, the fixed tissue samples were infiltrated with 2.3 M sucrose in PBS for 15 min. Frozen samples were sectioned at a thickness of 60 nm at −120°C and the sections transferred to formvar/carbon coated copper grids. Grids were floated on PBS, and downstream immunolabeling was carried out as above. Contrasting/embedding of the labeled grids was carried out on ice in 0.3% uranyl acetate in 2% methyl cellulose for 10 min.

### Size exclusion chromatography

Size exclusion chromatography of human brain soaking extracts was performed using ÄKTA FPLC and a Superdex 200 Increase 10/300 column (Cytiva). Extracts (350 μl) were injected into the column using a 0.5-ml loop. Running buffer was TBS (25 mM Tris, 150 mM NaCl, pH 7.4) at 0.5 ml/min. 0.5-ml fractions were collected and used directly for ELISA without lyophilization.

### Electron cryo-microscopy

Four to five 1-ml aliquots of soaking extracts were thawed on ice and spun in 1.5 ml Eppendorf tubes in an Eppendorf FA-45-24-11 rotor at 20,000 *g* for 1 h. All the pellets were resuspended and combined into 1 ml cold water. Following a 30 min centrifugation at 20,000 g at 4°C, pellets were resuspended in 10 μl of 20 mM Tris-HCl, pH 7.4. Samples were centrifuged at 3,000 *g* for 3 min and applied to glow-discharged holey carbon gold grids (Quantifoil Au R1.2/1.3, 300 mesh), which were glow-discharged with an Edwards (S150B) sputter coater at 30 mA for 30 s. Aliquots (3 μl) of samples were applied to the grids and blotted for approximately 3-5 s with filter paper (Whatman) at 100% humidity and 4 °C, using a Vitrobot Mark IV (Thermo Fisher). Datasets were acquired on a Titan Krios G3 microscope (Thermo Fisher Scientific) operated at 300 kV. Images for case AD7 were acquired using a Falcon-4 detector without energy filter. Images for case AD11 were acquired on a Gatan K3 detector in super-resolution counting mode, using a Bio-quantum energy filter (Gatan) with a slit width of 20 eV. Images were recorded with a total dose of 40 electrons per Å^2^.

Movie frames were gain-corrected, aligned, dose-weighted and then summed into a single micrograph using RELION’s own motion correction program.^41^ Contrast transfer function (CTF) parameters were estimated using CTFFIND-4.1.^42^ All subsequent image-processing steps were performed using helical reconstruction methods in RELION.^43,44^ Filaments were picked manually. Reference-free 2D classification was performed to select suitable segments for further processing. Deposited maps of Type I Aβ42 filaments (EMD-13800), Type II Aβ42 filaments (EMD-13809),^16^ and tau PHFs (EMD-3741 and EMD-0259)^16^ low-pass filtered to 10 Å were used as initial models. Combinations of 3D auto-refinement and 3D classifications were used to select the best segments for each structure. For all structures, Bayesian polishing^41^ and CTF refinement^43^ were performed to further increase the resolution of the reconstructions. Final reconstructions were sharpened using standard post-processing procedures in RELION, and overall resolutions were estimated from Fourier shell correlations at 0.143 between the independently refined half-maps, using phase randomisation to correct for convolution effects of a generous, soft-edged solvent mask.^45^

Atomic models comprising β-sheet rung were built in Coot,^46^ starting from available models for Type I Aβ42 filaments (PDB:7Q4B), Type II Aβ42 filaments (PDB:7Q4M), and PHFs (PDB:6HRE). Coordinate refinements were performed using *Servalcat*.^47^ Final models were obtained using refinement of only the asymmetric unit against the half-maps in *Servalcat*.

### Data availability

Cryo-EM maps have been deposited in the Electron Microscopy Data Bank (EMDB) under accession numbers EMD-15770, EMD-15771 and EMD-15772. Corresponding refined atomic models have been deposited in the Protein Data Bank (PDB) under accession numbers 8AZS, 8AZT and 8AZU.

