## Supplementary Tables for "Abundant Aβ fibrils in ultracentrifugal supernatants of aqueous extracts from Alzheimer’s disease brains"

| Brain | Age at death | Sex | *Postmortem* interval (h) | Other pathologic diagnoses |
| --- | --- | --- | --- | --- |
| AD1 | 73 | F | 20 | CAA, LBD of amygdala |
| AD2 | 79 | M | <36 | CAA |
| AD3 | 63 | F | <36 | Diffuse neocortical LBD |
| AD4 | 66 | F | <36 | CAA |
| AD5 | 65 | F | <48 | LBD of amygdala |
| AD6 | 73 | F | 10 | CAA |
| AD7 | 68 | F | 7 | Limbic LBD, CAA |
| AD8 | 75 | M | <36 | CAA |
| AD9 | 70 | F | <36 | Diffuse neocortical LBD, CAA |
| AD10 | 73 | M | 5 | Diffuse neocortical LBD, CAA |
| AD11 | 73 | M | <36 | None |
| AD12 | 68 | M | 65 | Down syndrome, CAA |
| AD13 | 83 | M | 2.5 | Diffuse neocortical LBD |
| C1 | 37 | F | <48 | None |
| C2 | 38 | M | <48 | None |
| C3 | 75 | M | <36 | Rare hippocampal neurofibrillary tangles. No amyloid pathology. |

**Table S1.** Cases of Alzheimer’s disease (AD) and neurologically normal controls (C).

CAA, cerebral amyloid angiopathy; LBD, Lewy body disease.

| Assay | Capture (concentration, source) | Detector (concentration, source) | Platform |
| --- | --- | --- | --- |
| Aβ_42_ monomer-preferring | m266 (3 µg/ml, Elan Pharmaceuticals) | 21F12 (0.4 µg/ml, Elan Pharmaceuticals) | MSD |
| Aβ_40_ monomer-preferring | m266 (3 µg/ml, Elan Pharmaceuticals) | 2G3 (0.2 µg/ml, Elan Pharmaceuticals) | MSD |
| Aβ aggregate-preferring | 71A1 (12.5 µg/mg beads, Abyssinia Biologics) | 3D6 (0.2 µg/ml, Elan Pharmaceuticals) | SMCxPRO |
| Aβ aggregate-preferring | 1C22 (12.5 µg/mg beads, gift of Dominic Walsh) | 3D6 (0.2 µg/ml, Elan Pharmaceuticals) | SMCxPRO |

**Table S2.** Sandwich ELISAs.

|  | **Type I Aβ** | **Type II** **Aβ** | **PHF** |
| --- | --- | --- | --- |
|  | (EMD-, PDB) | (EMD-, PDB) | (EMD-, PDB) |
| **Data acquisition** | | | |
| Electron gun | XFEG | | |
| Detector | K3 | | |
| Energy filter slit (eV) | 20 | | |
| Magnification | 105,000 | | |
| Voltage (kV) | 300 | | |
| Electron dose (e^–^/Å^2^) | 40 | | |
| Defocus range (μm) | -1.0 ~ -2.5 | | |
| Pixel size (Å) | 0.831 | | |
| **Map refinement** | | | |
| Symmetry imposed | C1 | C2 | C1 |
| Final particle images (no.) | 37380 | 14740 | 44535 |
| Map resolution (Å) | 2.9 | 3.7 | 3.1 |
| FSC threshold | 0.143 | 0.143 | 0.143 |
| Helical twist (°) | 178.4 | -2.8 | 179.4 |
| Helical rise (Å) | 2.4 | 4.8 | 2.4 |
| **Model refinement** |  |  |  |
| Model resolution (Å) | 2.9 | 3.7 | 3.1 |
| FSC threshold | 0.5 | 0.5 | 0.5 |
| Map sharpening *B* factor (Å^2^) | -64 | -103 | -79 |
| Model composition |  |  |  |
| Non-hydrogen atoms | 1506 | 904 | 3976 |
| Protein residues | 204 | 124 | 518 |
| Ligands | 0 | 0 | 0 |
| *B* factors (Å^2^) |  |  |  |
| Protein | 81.1 | 77.3 | 76.1 |
| R.m.s. deviations |  |  |  |
| Bond lengths (Å) | 0.0078 | 0.0067 | 0.0071 |
| Bond angles (°) | 1.36 | 1.36 | 1.47 |
| Validation |  |  |  |
| MolProbity score | 1.31 | 1.44 | 2.28 |
| Clashscore | 3.31 | 2.17 | 4.80 |
| Poor rotamers (%) | 0 | 0 | 0 |
| Ramachandran plot |  |  |  |
| Favored (%) | 96.88 | 93.10 | 84.72 |
| Allowed (%) | 3.12 | 6.90 | 15.28 |
| Disallowed (%) | 0 | 0 | 0 |

**Table S3.** Cryo-EM data acquisition and structure determination.
